# A highly conserved two-gene operon is crucial for lipoarabinomannan localization, pathogenesis, and cell envelope function in *Mycobacterium abscessus*

**DOI:** 10.64898/2026.01.29.702501

**Authors:** Nicholas Campbell-Kruger, Amir Balakhmet, Sarah A. Stanley

## Abstract

*Mycobacterium abscessus* is an emerging threat, causing infections that are difficult to treat due to intrinsic resistance to most antibiotics. Determinants of *M. abscessus* physiology and pathogenesis remain poorly understood, hampering therapeutic development. Here, we show that in *M. abscessus*, the *lprg-mfs* operon is essential for virulence in macrophages and in mice. Loss of *lprg-mfs* in *M. abscessus* causes accumulation of the glycolipid lipoarabinomannan (LAM) on the cell surface and in culture supernatant suggesting that this system participates in LAM import. This contrasts with its proposed role in *M. tuberculosis* where *lprg-mfs* has been implicated in the export of various lipids. Consistent with altered lipid distribution, the *lprg-mfs* mutant displays severe defects in mycomembrane permeability, fluidity, and integrity, and expression of *mfs* alone restores only a subset of these phenotypes, revealing a surprising uncoupling of envelope fluidity and permeability. Using a suppressor screen to further investigate factors that control the distribution of lipoarabinomannan we find that a point mutation in the unannotated gene *MAB_0995* can fully or partially complement all deletion mutant phenotypes. Our data also show that lipoarabinomannan in the mycomembrane is dynamically regulated in response to environmental conditions, including hypoxia and macrophage infection. Together, these findings redefine the role of LprG/Mfs in mycobacterial cell envelope homeostasis and reveal unexpected plasticity in mycomembrane lipid regulation in *M. abscessus*.

**Importance:** The emerging pathogen *Mycobacterium abscessus* causes life-threatening lung infections in certain patients that are extremely difficult to treat due to its intrinsic resistance to most antibiotics. However, the process by which this organism establishes infection is poorly understood, as are the specific determinants of antibiotic tolerance. Better knowledge of the genes required for virulence and impermeability to antibiotics in *M. abscessus* could enable to development of more effective treatments. The significance of this study is the demonstration that the *lprg-mfs* operon is required both for pathogenesis and for impermeability in *M. abscessus*. Further, our study shows a correlation between cell envelope characteristics and the distribution of the molecule lipoarabinomannan, suggesting a specific mechanism by which these crucial characteristics are mediated.

## Introduction

The Gram-positive family *Mycobacteriaceae* contains many important human and animal pathogens, including *Mycobacterium tuberculosis*, a pathogen responsible for over one million deaths annually (1). A defining feature of mycobacteria is the mycomembrane, a secondary membrane external to the peptidoglycan that contributes both to pathogenesis and antibiotic resistance. The inner leaflet of the mycomembrane consists of extremely long chain, branched carboxylic acids called mycolic acids covalently connected to the peptidoglycan via a branched polysaccharide arabinogalactan. The outer leaflet is more chemically diverse and contains trehalose mono- and di-mycolates as well as a variety of lipids that are crucial for pathogenesis, including the highly glycosylated phosphatidylinositol lipoarabinomannan (LAM) (2–4). While the biosynthetic pathways of these lipids are largely elucidated, the mechanisms by which these lipids are transported through the aqueous periplasm and inserted into the mycomembrane, and the extent to which mycobacteria dynamically regulate and maintain this membrane, are still unknown.

*Mycobacterium abscessus* is an emerging opportunistic pathogen with a globally increasing disease burden (5). *M. abscessus* is intrinsically resistant to most antibiotics, and infections have low cure rates of ∼50% (6). The bacterial factors that enable *M. abscessus* to establish infection, survive a variety of host stressors, and exhibit innate resistance to antibiotic therapies are poorly understood. As the outer layer of the cell, the mycomembrane is likely to be essential in some or all these processes, but its composition is poorly characterized in *M. abscessus* compared to other, better-studied mycobacteria (7). In addition, the fundamental processes of lipid transport, insertion, and regulation are still understudied in *M. abscessus*.

The *lprg-mfs* operon consists of the lipoprotein LprG (MAB_2806) and a Major Facilitator Superfamily transporter (MAB_2807) and is highly conserved across mycobacteria. This operon is crucial for virulence in *M. tuberculosis*, but the underlying mechanisms are unclear (8, 9). *In vitro*, LprG from *M. tuberculosis* binds to a variety of triacylated lipids, including triacylglycerides, phosphatidyl-inositol mannosides, and the more highly glycosylated derivatives lipomannan and LAM (8, 10, 11). Additionally, work in non-tuberculous mycobacteria has shown that *lprg-mfs* mutants exhibit a pronounced sensitivity to a broad spectrum of antibiotics, likely caused by a membrane stiffness-mediated permeability defect (12, 13). The current model of *lprg-mfs* function is that this system exports components of the mycomembrane which are critical for impermeability and pathogenesis, but important assumptions of this model remain hypothetical. It is unclear which of the reported substrates of *lprg-mfs* are important in a physiological context, how these substrates contribute to the observed permeability and virulence defects, and whether this operon performs the same role in different mycobacterial species.

Here we show that in contrast to mutants in *Mtb*, loss of *lprg-mfs* results in increased, rather than decreased, abundance of LAM in the outer layer of *M. abscessus*. Loss of this operon has profound impacts on mycomembrane permeability, stiffness, and integrity. Furthermore, we show that a *lprg-mfs* deletion mutant has attenuated virulence. Interestingly, we show that complementation of *mfs* alone is sufficient to restore some, but not all, phenotypes, suggesting nonredundant roles for *lprg* and *mfs*. We also show that LAM localization, and more broadly mycomembrane characteristics, are partially or fully restored in a *lprg-mfs* deletion mutant by a point mutation in the uncharacterized gene *MAB_0995*. Finally, we show for the first time that LAM localization in *M. abscessus* changes dynamically in response to different environmental conditions. Our findings suggest that LAM content in and beyond the mycomembrane is a key factor determining cellular characteristics, and that *lprg-mfs* activity determines LAM content of these cellular compartments.

## Results

### *lprg-mfs* is important for pathogenesis in *M. abscessus*

To determine whether the *lprg-mfs* operon contributes to pathogenesis in *M. abscessus*, we generated a mutant lacking this operon (hereafter referred to as dKO) using ORBIT (14) as well as a complemented strain using an integrative plasmid containing *lprg-mfs* under the control of its native promoter. To avoid potential interactions with the known virulence determinant glycopeptidolipid (GPL), these strains were generated in a smooth background. Murine bone marrow-derived macrophages (BMMs) infected with WT, dKO, and complemented strains of *M. abscessus*. We observed that WT and complemented *M. abscessus* increases in number over the first three days of infection. In contrast, dKO remains static, leading to a seven-fold difference in bacterial number on day three of infection (Figure 1A). For all strains, no reduction in BMM viability is observed when compared to uninfected BMMs, demonstrating that differences in bacterial burdens between strains do not reflect differences in macrophage survival (Figure 1B).

**Figure 1.**
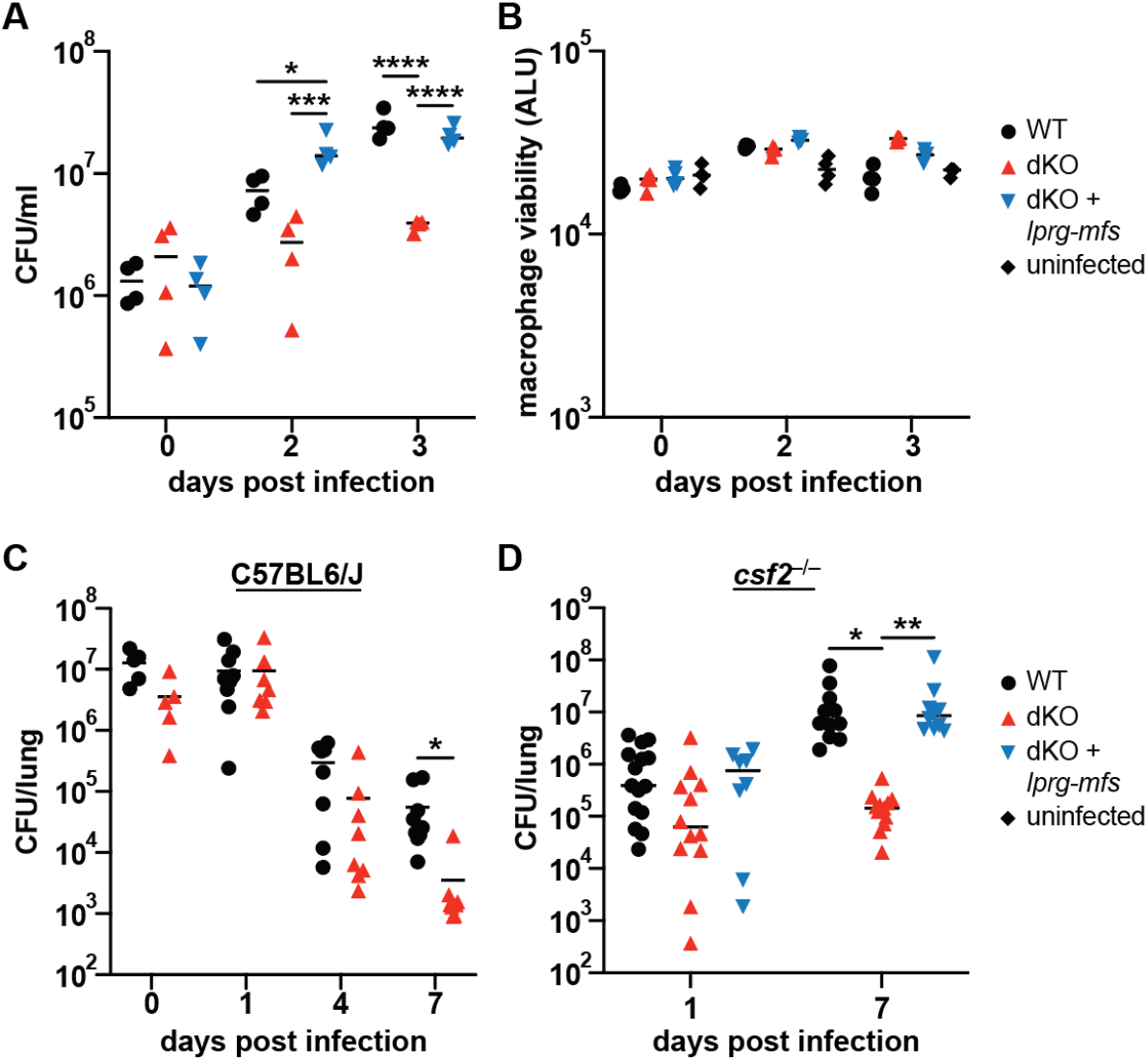
Impact of *lprg-mfs* deletion on virulence in *M. abscessus*. **A**,**B**: Murine BMMs were infected with WT, dKO, and complemented *M. abscessus* and **(A)** CFU or **(B)** macrophage viability via CellTiter Glo 2.0 were monitored at indicated timepoints. **(C)** C57BL6/J mice were infected i.n. with 10^7^ *M. abscessus* cfu and bacterial burden was enumerated in whole lungs. Data are pooled from two experiments. (**D)** *Csf2*^*–/–*^ mice were infected i.n. with 10^5^ *M. abscessus* CFU and bacterial burden was enumerated in whole lungs. Data are pooled from three experiments. * p≤.05, ** p≤ .001, *** p≤.0005, **** p≤.0001. **(A)** Two-way ANOVA with Tukey multiple comparison correction, (**C**) Mann-Whitney U test. **(D)** Two-way ANOVA with Tukey multiple comparison correction.

C57BL/6J and other immunocompetent mouse strains are known to effectively control infection with wild type *M. abscessus* (15). Indeed, intranasal (i.n.) infection of C57BL/6J mice with 10^7^ CFU of wild type *M. abscessus* leads to a hundred-fold reduction in bacterial burden over the first seven days (Figure 1C). Mice infected with dKO *M. abscessus* exhibit more rapid control, with a tenfold reduction in bacterial burden compared to WT *M. abscessus* at this time point (Figure 1C). The limitation of rapid clearing in wild type infections in mice has led to the generation of a wide variety of more permissive animal models for *M. abscessus* infection including mice lacking GM-CSF (*Csf2*^*–/–*^) (16, 17). In contrast with C57BL/6J mice, *Csf2*^*–/–*^ mice are unable to control intranasal infection of WT *M. abscessus* (Figure 1D). dKO bacterial burden does not increase over the same time frame, resulting in a hundred-fold difference in bacterial burden at day 7 (Figure 1D). Thus, in both immunocompetent and immunocompromised models of infection, *lprg-mfs* is important for virulence of *M. abscessus*.

### Deletion of *lprg-mfs* leads to altered mycomembrane attributes in *M. abscessus*

Production of GPL, a component of the mycomembrane, plays a role in modulating virulence of *M. abscessus* (18, 19). Loss of GPL induces the transition from smooth to rough, a phenotype that is assayed by colony morphology (20). Interestingly, dKO *M. abscessus* adopts a distinct morphology when grown on multiple media formulations (Figure 2A). This colony morphology is different from the smooth to rough transition, as dKO *M. abscessus* does not have altered levels of GPL when compared to WT (Figure S1A). Given the differences observed in colony morphology, we next addressed whether the mycomembrane in dKO *M. abscessus* is functionally impaired. As previously shown in *M. smegmatis*, dKO *M. abscessus* exhibits dramatically increased permeability to ethidium bromide (Figure 2B). dKO *M. abscessus* also exhibits increased membrane stiffness when measured via Laurdan (6-dodecanoyl-2-dimethylaminonaphthalene) (Figure 2C). As expected for a bacterial strain with altered envelope integrity, dKO *M. abscessus* is markedly more susceptible to both cationic and anionic detergents (Figure 2D,E). Thus, deletion of *lprg-mfs* dramatically changes the permeability, fluidity, and integrity of the mycobacterial cell envelope, and alters the colony morphology of *M. abscessus*.

**Figure 2.**
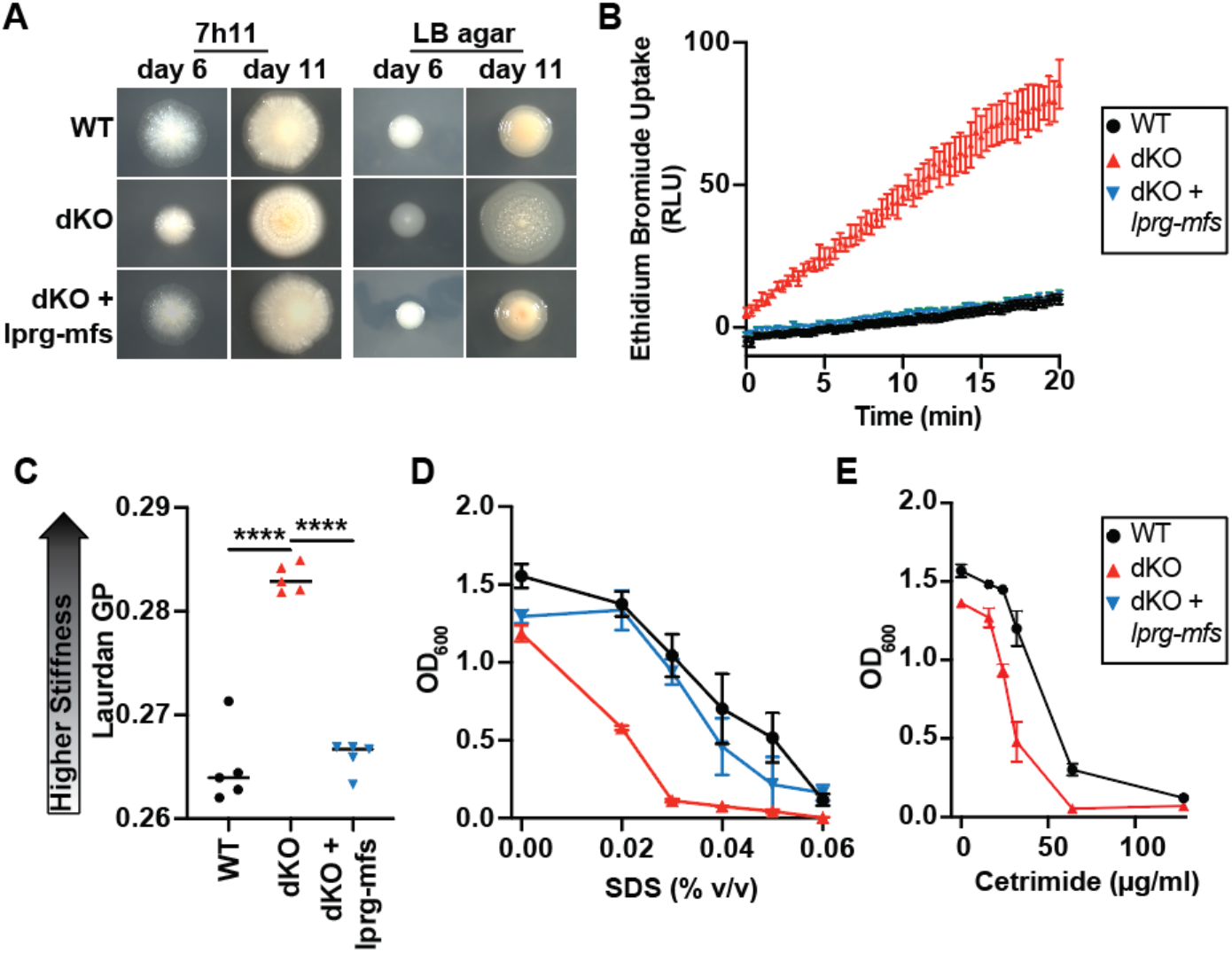
Impact of *lprg-mfs* deletion on colony morphology and mycomembrane characteristics. (**A**) WT, dKO, and complemented *M. abscessus* colonies imaged with a stereomicroscope. (**B**) Strains were cultured in 7H9 and mycomembrane permeability was measured via EtBr uptake. (**C**) Strains were cultured in 7H9 and mycomembrane fluidity was measured via Laurdan generalized polarity. **D**,**E:** Strains were cultured in 7H9 in the presence of SDS **(D)** or cetrimide **(E). (B**,**D**,**E)** data plotted are mean ± standard deviation. **** p≤.0001 by **(C)** Ordinary one-way ANOVA with Tukey’s post-hoc test.

### Deletion of *lprg-mfs* leads to upregulated lipoarabinomannan export in *M. abscessus*

LprG has been suggested to bind to and export numerous lipids, including TAG, PIM and LAM. We used a reverse micelle extraction to specifically extract lipids in the outer leaflet of the mycomembrane (4) and visualized these fractions through TLC. A variety of expected lipids were recovered, but no PIMs were detected in this fraction and no differences in the abundance of TAGs could be found (Figure S1B,C). We next assayed cell supernatants and cellular lysates for LAM using the CS-35 monoclonal antibody that has been shown to bind robustly to LAM from multiple mycobacterial species (21–23). Surprisingly, dKO *M. abscessus* exhibits markedly increased LAM in the culture supernatant (Figure 3A). No differences in LAM abundance could be found in whole cell lysates, indicating that the increased abundance of LAM in culture supernatant of dKO *M. abscessus* is due to differential localization of LAM rather than a difference in gross abundance (Figure 3A).

**Figure 3.**
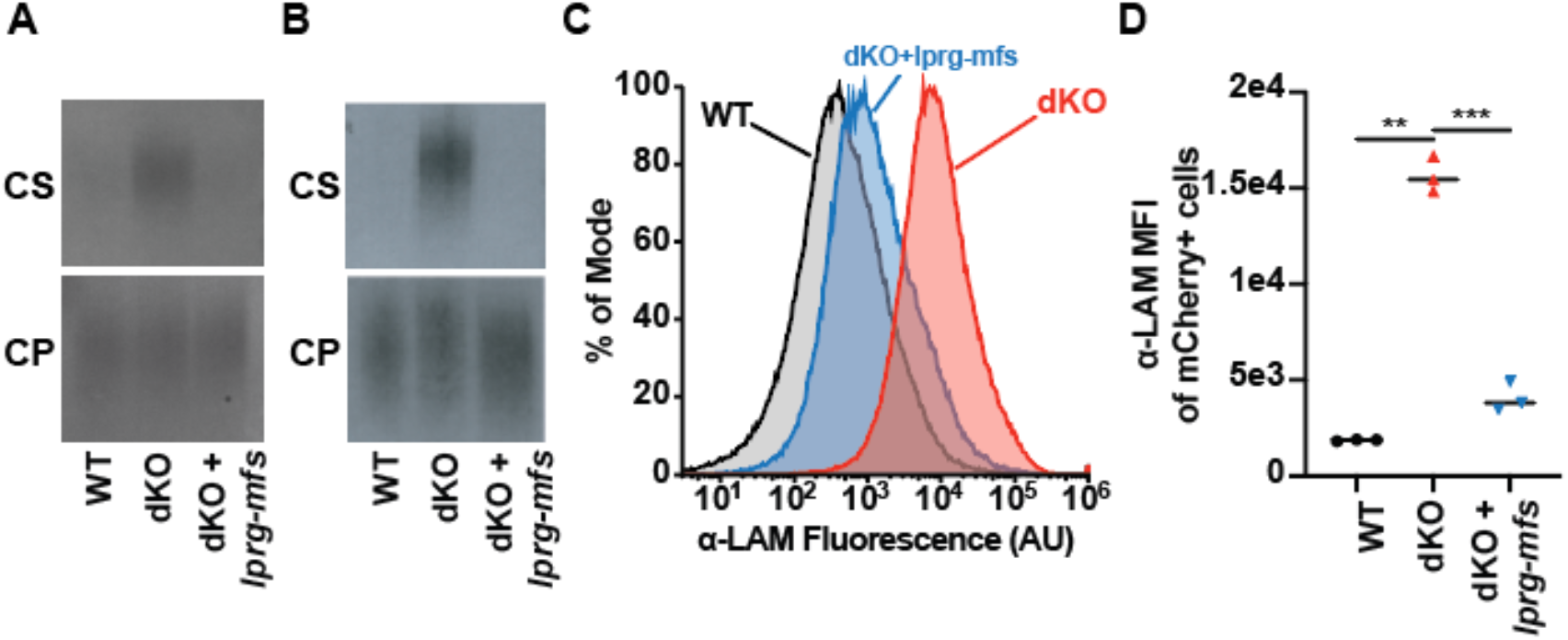
Impact of *lprg-mfs* deletion on lipoarabinomannan content and localization. **A**,**B:** Western blots of culture supernatant (CS) and cell pellet (CP) lipoarabinomannan content. Cells were cultured in 7H9 **(A)** or BSA-free media **(B)** and analyzed using the α-LAM antibody CS-35. **C**,**D:** whole, unpermeabilized cells were fixed, stained for surface exposed LAM, and analyzed by flow cytometry. Shown are representative flow plots **(C)** and quantification of three replicates **(D)** of the α-LAM mean fluorescence intensity of mCherry positive single cells. ** p≤ .001, *** p≤.0005. Brown-Forsythe ANOVA.

Traditional 7H9 medium contains bovine serum albumin, a promiscuous lipid-binding protein. Albumin might increase the amount of LAM in the culture supernatant by binding to the lipidic moiety of the molecule, extracting it from the lower integrity mycomembrane of dKO *M. abscessus*. We cultured *M. abscessus* in 7H9 supplemented with additional glycerol and dextrose rather than OADC to remove this variable while minimizing compositional changes. Under these conditions LAM is completely absent in the culture supernatant of WT *M. abscessus*, while dKO *M. abscessus* still sheds significant amounts of LAM despite no differences in whole cell abundance (Figure 3B).

Differences in abundance in culture supernatant may not be reflective of compositional changes of the mycomembrane, so we next quantified LAM located on the cell surface using flow cytometry of whole, unpermeablized bacterial cells. Fluorescent strains of *M. abscessus* were generated by transforming existing strains with an episomal plasmid containing *mcherry* under the control of a constitutive promoter to differentiate bacterial cells from debris. In agreement with the analysis of culture supernatant, dKO *M. abscessus* exhibits significantly higher amounts of surface-exposed LAM (Figure 3C,D). Previous work in *M. tuberculosis* has shown that deletion of *lprg* alone leads to decreased, rather than increased, LAM on the cell surface (11, 24). In contrast to these reports, deletion of *lprg-mfs* leads to a dramatic increase in the amount of LAM found on the cell surface and in the culture supernatant of *M. abscessus*.

### *mfs* alone is sufficient for complementation of some, but not all, dKO phenotypes

Understanding the contributions of LprG and Mfs individually might shed light on the function of the LprG-Mfs system, especially considering possible crosstalk with other lipid transport systems. We generated two additional complementation strains by transforming dKO *M. abscessus* with integrative plasmids containing *lprg* or *mfs*, each under the control of the native *lprg-mfs* promoter. In line with work in *M. smegmatis* (12), *mfs* is sufficient to restore WT-like ethidium bromide permeability (Figure 4A). Interestingly, *mfs* alone can also restore SDS sensitivity, but not membrane stiffness (Figure 4B,C). *Mfs* complements the increase in LAM observed in culture supernatant of the dKO mutant but does not complement the increase in surface-exposed LAM by flow cytometry (Figure 4D-F). Complementation of *lprg* alone is unable to restore any of the phenotypes interrogated in this report (Figure 4A-F). These results show that Mfs has an LprG independent role in regulating the characteristics of the *M. abscessus* cell envelope.

**Figure 4.**
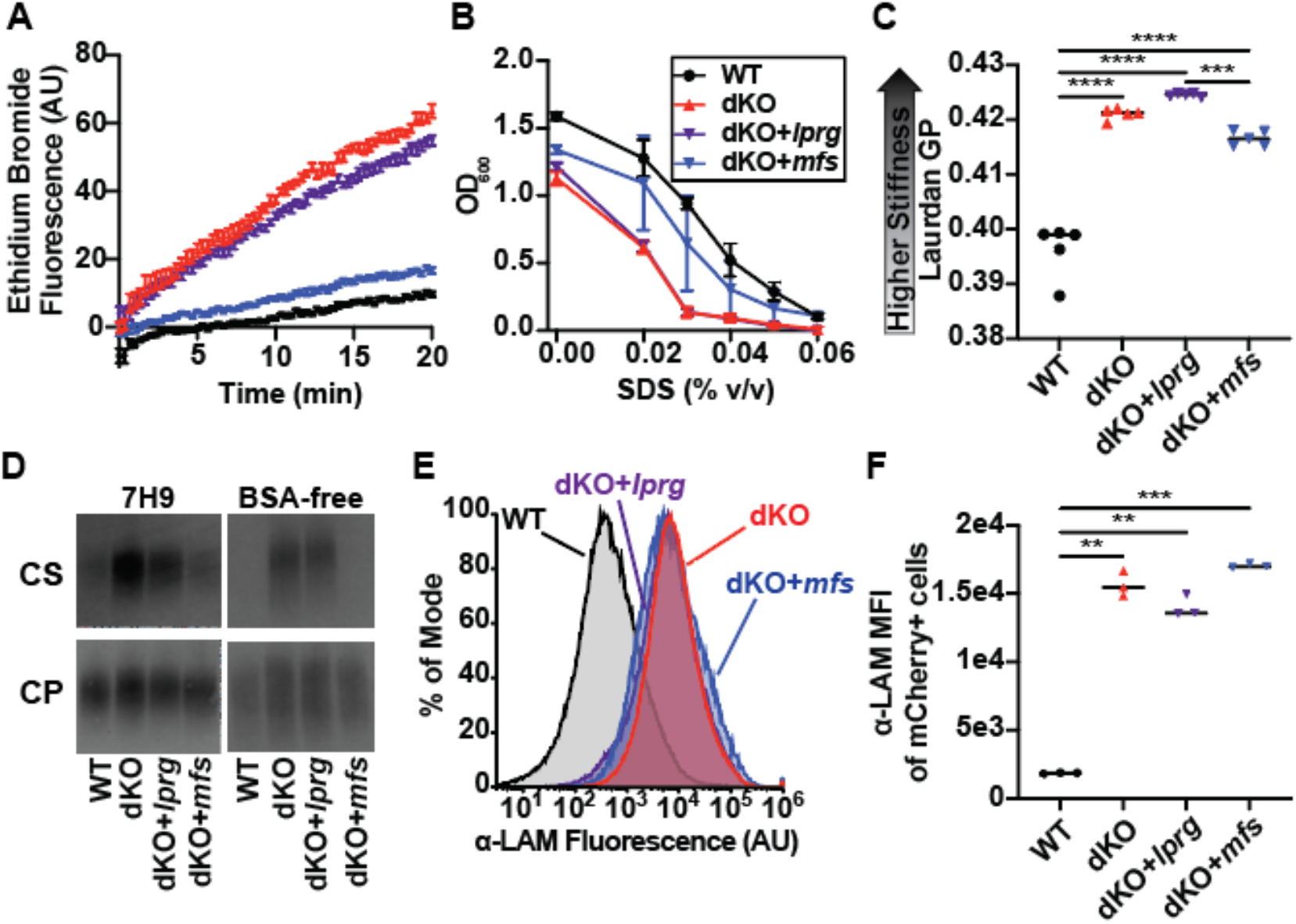
Impact of single gene complementation on dKO phenotypes. **(A)** Membrane permeability of single gene complementation strains measured through ethidium bromide uptake. **(B)** SDS sensitivity of single gene complementation strains. **(C)** Membrane fluidity of single gene complementation strains. **(D)** Western blot of LAM content of single gene complementation strain culture supernatants (CS) and cell pellets (CP) in 7H9 and BSA-free media. **E**,**F:** Whole, unpermeabilized cells were analyzed by flow cytometry. Representative flow plots **(E)** and quantification of 3 independent replicates **(F)**. ** p≤ .001, *** p≤.0005, **** p≤.0001. **(C)** Ordinary one-way ANOVA. **(F)** Brown-Forsythe ANOVA

### A point mutation in an uncharacterized gene largely restores dKO *M. abscessus* to WT phenotypes

To identify genes that impact LAM-dependent integrity of the mycomembrane, we performed a suppressor screen for mutations that restore dKO *M. abscessus* to WT-like detergent sensitivity. dKO *M. abscessus* was cultured in 7H9 with no detergent and two billion cells were plated on 7H10 plates containing 0.03% SDS. Four resultant colonies (Sup1-Sup4) were subjected to an in-broth assay to confirm SDS resistance, and the genomes of all four colonies were sequenced (Figure 5A). All were found to have a single mutation in common when compared to the parental dKO genome. This mutation, a T->G substitution, is a missense W89G mutation in the uncharacterized gene *MAB_0995*, a putative oxidoreductase.

**Figure 5.**
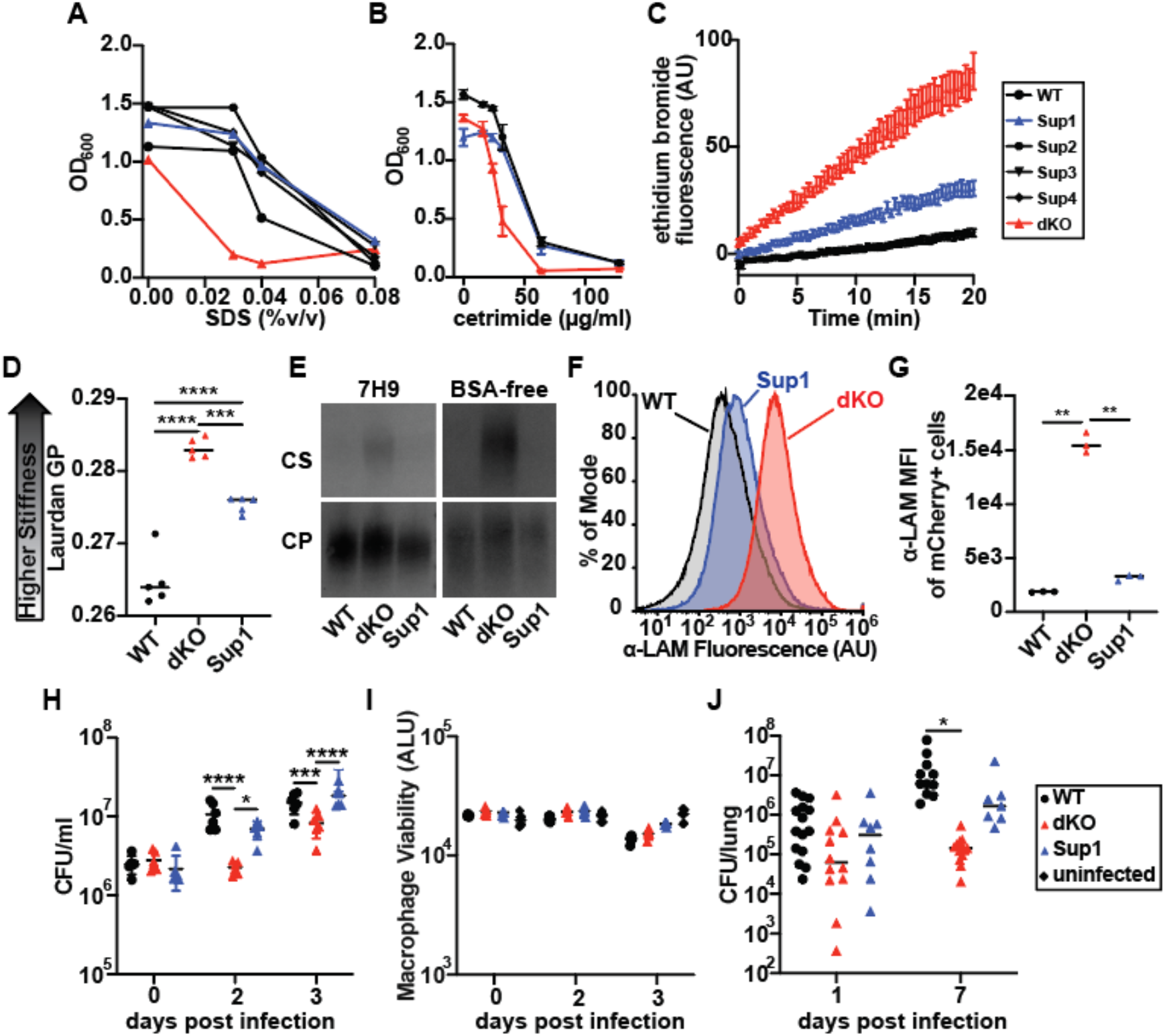
Effect of SDS suppressor mutant Sup1 on dKO phenotypes. **(A)** Confirmation of SDS resistance in putative suppressor mutants. **(B)** Strains were cultured in 7H9 in the presence of cetrimide. **(C)** Membrane permeability of Sup1 measured via ethidium bromide uptake. **(D)** Membrane fluidity of Sup1 measured through Laurdan GP. **E-G:** Quantification of LAM via western blot of culture supernatant (CS) and cell pellet (CP) **(E)** or flow cytometry **(F-G). H**,**I:** Murine BMMs were infected with Sup1 and assessed via bacterial enumeration **(H)** and CellTiter Glo 2.0 **(I). (J)** *Csf2*^−/−^ mice were infected intranasally with Sup1 and bacterial number was assessed after 7 days. * p≤.05, ** p≤ .001, *** p≤.0005, **** p≤.0001. **(G)** Brown-Forsythe ANOVA. **(H)** Ordinary two-way ANOVA. **(J)** Two-way ANOVA.

As all colonies isolated contained the same mutation, we further characterized the impact of this mutation using the suppressor mutant Sup1. To test whether Sup1 impacts mycomembrane characteristics beyond anionic detergent sensitivity, we assayed cationic detergent sensitivity, membrane permeability via ethidium bromide uptake and membrane fluidity via Laurdan. Sup1 has WT-like cetrimide sensitivity and partially restores membrane permeability and fluidity (Figure 5B-D). Having found that Sup1 appears to broadly suppress dKO phenotypes, we interrogated LAM content through western blot and flow cytometry. Sup1 completely suppresses the high levels of LAM found in dKO culture supernatant and on the cell surface (Figure 5E-G). To assay the virulence of Sup1, we infected BMMs and monitored bacterial number and BMM viability over time. Sup1 replicates in BMMs to the same degree as WT *M. abscessus*, overcoming the dKO defect in virulence (Figure 5H). As seen previously, none of the tested strains caused a significant decrease in BMM viability compared to uninfected controls (Figure 5I). Having found that Sup1 is sufficient to restore virulence *in vitro*, we next tested this strain *in vivo*. When *Csf2*^*–/–*^ mice were infected intranasally with Sup1, intermediate levels of bacteria between WT and dKO *M. abscessus* were recovered after seven days (Figure 5J). Although not statistically significant, these data suggest that Sup1 partially restores WT-like virulence *in vivo*.

### LAM content of the mycomembrane is dynamically regulated in *M. abscessus*

Previous work on *lprg-mfs* has largely concluded that this operon functions to export lipid species to the mycomembrane or beyond the boundaries of the cell. The surprising result that deletion of this operon in *M. abscessus* leads to an increase, rather than decrease, of LAM export suggests that *lprg-mfs* could function instead as a retrograde transporter of LAM or that *lprg-mfs* transport activity could be bidirectional and controlled through other yet-unknown factors. If the latter model is true, environmental stimuli might control the direction of transport and lead to differential dKO phenotypes. To test this model, we assayed the LAM content of the mycomembrane in response to hypoxia, a physiologically relevant condition that induces significant changes in the metabolism and cell envelope structure of mycobacteria (25–27). *M. abscessus* cultures were cultured in sealed glass tubes to prevent gas exchange and methylene blue was used to monitor the oxygenation state of the cultures. All cultures decolorized completely within three days, at which point the cultures were analyzed by flow cytometry. All tested strains of *M. abscessus* significantly upregulate surface-exposed LAM under hypoxia (Figure 6A,B). WT *M. abscessus* technically exhibits a greater-fold change induced by hypoxia due to the lower starting amount of LAM, but dKO *M. abscessus* upregulated LAM to equivalent levels as WT, suggesting that *lprg-mfs* is not required to increase LAM levels on the cell surface.

**Figure 6.**
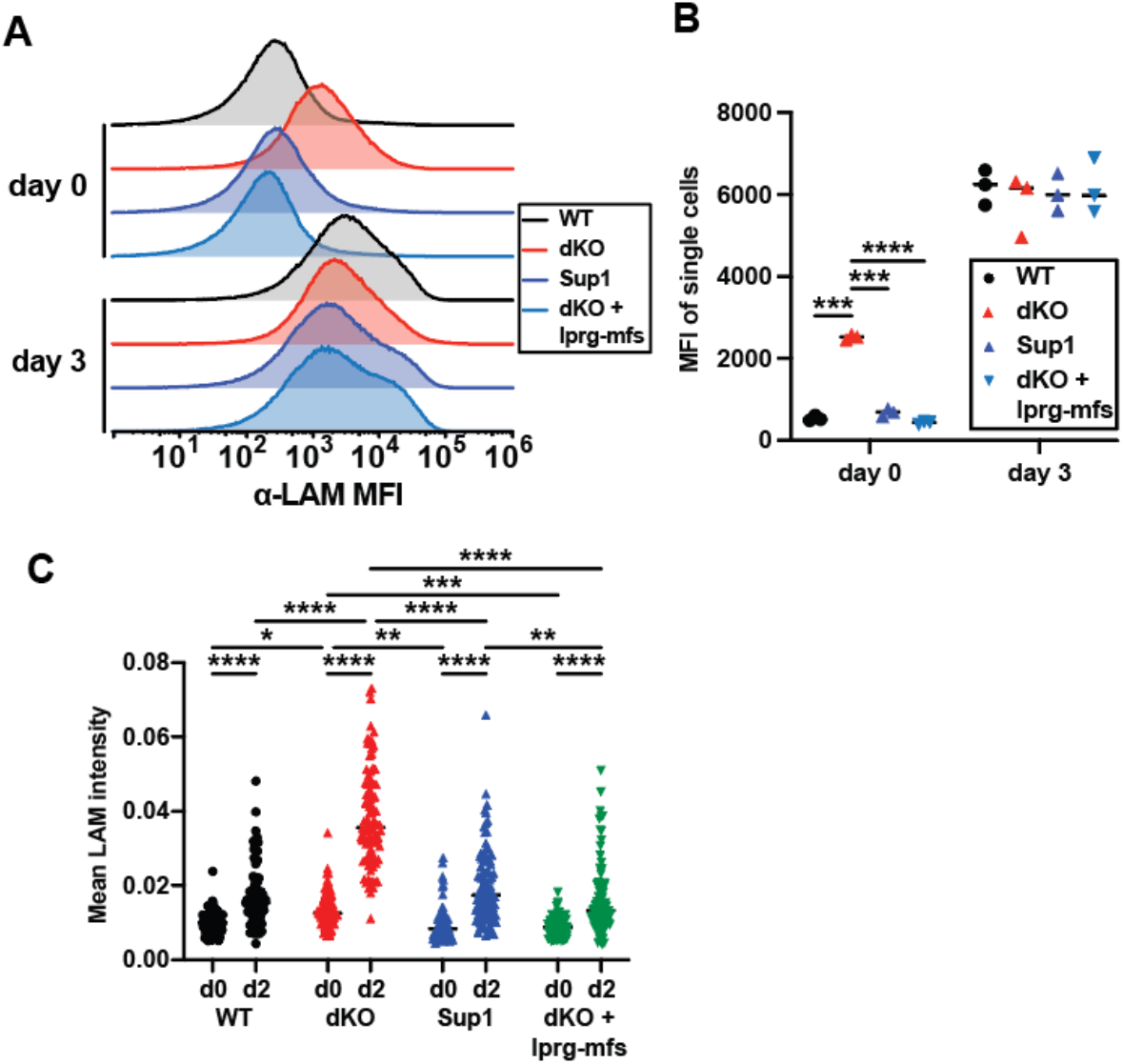
Hypoxia and macrophage infection cause changes in LAM abundance of *M. abscessus*. **A**,**B:** Quantification of LAM by flow cytometry for *M. abscessus* before and after hypoxia. Representative flow plots **(A)** and quantification of three independent replicates **(B). (C)** Quantification of LAM signal localized to mCherry+ bacteria by fluorescence microscopy of *M. abscessus* in *in vitro* macrophage infection. * p≤.05, ** p≤ .001, *** p≤.0005, **** p≤.0001. **(B)** Brown-Forsythe ANOVA. **(C)** Two-way ANOVA

Having found that *M. abscessus* LAM content in the mycomembrane changes under hypoxia, we wondered whether a similar modulation might occur in the context of a macrophage infection. BMMs were infected with fluorescent *M. abscessus*, immunofluorescence was used to label LAM, and images were analyzed through a CellProfiler pipeline. All strains of *M. abscessus* significantly upregulate LAM through the course of infection (Figure 6C). As was seen in hypoxia, *M. abscessus* upregulates the LAM content of its mycomembrane through the course of BMM infection in a process that appears independent of *lprg-mfs*.

## Discussion

*M. abscessus* is an important emerging pathogen, but the determinants of pathogenesis are poorly understood in this bacterium. *M. abscessus* lacks many of the crucial determinants of virulence in *M. tuberculosis*, including the ESX-1 secretion apparatus and the related lipids phthiocerol dimycocerosate and phenolic glycolipid. In this report, we show that the *lprg-mfs* operon is crucial for *M. abscessus* pathogenesis both *in vitro* and *in vivo*. LAM has long been known to be important in determining the immune response in the context of *M. tuberculosis* infection (28–30), and more work elucidating the role of this molecule in *M. abscessus* virulence may shed further light on the pathogenesis of this poorly understood organism.

Previous work supports the idea that both *lprg* and *mfs* are lipid- or LAM-binding proteins, and that deletion of this operon has profound effects on the permeability of the mycomembrane to a variety of antimicrobial compounds. Additionally, *lprg* mutants have been found to have decreased LAM on the cell surface of *M. tuberculosis* (11, 24). These findings support a model in which *lprg-mfs* functions as an anterograde transporter of LAM, exporting it to the mycomembrane or into the capsule and supernatant. Here we report the opposite result: LAM abundance is sharply increased on the surface of the cell and in the culture supernatant of dKO *M. abscessus*. If *lprg-mfs* directly transports LAM, it must function as a retrograde rather than anterograde transporter. Alternatively, if *lprg-mfs* is an anterograde transporter, its substrate may be another lipid, and LAM overabundance may be compensatory. We found no differences in the composition of the dKO mycomembrane in this study using TLC analysis, suggesting that *lprg-mfs* is a transporter of LAM in *M. abscessus* as it is in *M. tuberculosis*, but that the direction of transport is inverted. This difference in function could be explained by the relatively large divergence between *M. tuberculosis* and *M. abscessus* (31). The *lprg-mfs* operon is strictly conserved within mycobacteria, but the divergence between these two species is seen at the level of LprG protein sequence, and it is feasible to conclude that this operon has evolved to distinct functions in these two species (12). However, it has been shown that both *M. abscessus* and *M. tuberculosis* operons can complement a deletion mutant in *M. smegmatis*, so more work detailing the differences in function between mycobacterial species is needed to shed light on the direction of transport in this system.

It is intriguing that expression of *mfs* alone is sufficient to complement detergent sensitivity, ethidium bromide uptake, and high levels of LAM found in culture supernatant, but the complete operon is required to restore membrane fluidity and reduce the amount of LAM found on the cell surface. One possibility is that *mfs* might determine the LAM content of both the mycomembrane and the capsule, while *lprg* is required for only one of these compartments. In this model, the requirement for both genes to restore both cell surface LAM and membrane stiffness implies that *lprg* helps to modulate mycomembrane LAM content, and that mycomembrane LAM content is crucial for controlling the stiffness of this membrane. Intriguingly, recent work has shown that mycobacteria modulate inner membrane stiffness through the further acylation of phosphatidylinositol mannosides already present in that compartment, potentially playing a role in *in vivo* antibiotic tolerance (32, 33). The lipidic component of LAM is phosphatidylinositol mannoside, suggesting a mechanism by which the increased abundance of LAM in the mycomembrane might cause the observed increase in stiffness of the dKO mutant. Further work examining the acylation state of LAM and interrogating how the composition of the mycomembrane might change in response to various stressors could further elucidate the putative link between membrane stiffness and LAM in the mycomembrane.

Our observed separation between membrane stiffness and membrane permeability phenotypes is surprising. Conditions leading to a stiffened envelope in *E. coli* dramatically reduce antibiotic resistance, likely through a permeability defect, and in general there is an established relationship between membrane fluidity and permeability or integrity in a variety of bacteria (34). It is thus unclear how *mfs* alone can complement membrane permeability in dKO *M. abscessus* without also complementing membrane stiffness. One possibility is that ethidium and Laurdan both report on the average characteristics of the whole cell rather than those of any particular membrane. We cannot rule out the possibility that the measured changes in stiffness and permeability are caused by perturbations to different cellular compartments.

We conducted a suppressor screen against the SDS sensitivity observed in dKO *M. abscessus*. Sup1, a strain of dKO *M. abscessus* bearing a single base substitution in the uncharacterized gene *MAB_0995*, exhibits WT-like sensitivity to detergents. Indeed, Sup1 generally suppresses dKO phenotypes, including LAM localization and virulence. Interestingly, while Sup1 fully restores WT-like virulence in a macrophage infection model, it only partially suppresses the dKO virulence defect *in vivo*. Similarly, Sup1 almost completely restores WT-like LAM in the mycomembrane and supernatant, but only partially restores membrane permeability and fluidity.

Taken together, these data suggest that the virulence defect observed *in vitro* is largely determined by LAM localization, but that membrane fluidity and permeability might be important determinants of *in vivo* pathogenesis. MAB*_0995* is predicted to encode an F_420_-dependent monooxygenase, and shares features specifically with dehydrogenases (35, 36). Despite this classification, the specific functions and substrates of MAB_0995 remain unknown. Recent work has found that in a lung infection model, *M. abscessus* requires biotin synthesis to properly modulate envelope fluidity through the production of unsaturated and branched-chain fatty acids, lending support to the idea that membrane fluidity may be important under physiological conditions (37). Future work dissecting the relationship between membrane fluidity or permeability and *in vivo* pathogenesis could reveal important aspects of the biology of *M. abscessus*.

In summary, we find that the conserved *lprg-mfs* operon is crucial for virulence and mycomembrane function in the emerging pathogen *M. abscessus*. We show the surprising result that in a strain lacking this operon, LAM is sharply increased in both the mycomembrane and culture supernatant and shed light on the different roles played by *lprg* and *mfs* individually. We find that LAM content in these compartments can be controlled by multiple other factors, such as a mutation in the uncharacterized gene *MAB_0995* and different environmental conditions. Our results give important insight into the poorly understand pathogenesis of *M. abscessus* and suggest a crucial link between LAM distribution and cell envelope integrity, permeability, and stiffness.

## Materials and Methods

### Bacterial strains and media

*M. abscessus* strain ATCC19977 was grown in Middlebrook 7H9 supplemented with 10% OADC, .05% Tween-80, and .2% glycerol. For experiments in which culture supernatant was assayed for LAM content, Tween-80 was omitted from the media. When growth on solid media was required, *M. abscessus* was grown on LB agar plates for 5-7 days. When necessary, kanamycin and zeocin were added to media at concentrations of 50μg/ml and 75μg/ml respectively, and hygromycin was added at concentrations of 100μg/ml in liquid media and 1mg/ml in solid media.

### Genetic manipulation of M. abscessus

To generate dKO *M. abscessus*, ORBIT was performed as previously described (14). Briefly, an ORBIT competent strain of *M. abscessus* was created via transformation with pKM444. This strain was grown to an OD_600_ 0.1-0.6, induced with anhydrotetracycline for 4 hours, and made electrocompetent as described (38). Cells were transformed with the zeocin-resistant payload vector pKM496 and a targeting oligonucleotide. After proper insertion was confirmed via PCR and sequencing, cells were cured of pKM444 via parallel streaking on LB with and without kanamycin.

### In vitro infections

Bone marrow derived macrophages (BMMs) isolated from C57BL/6J mice were seeded at 50,000 cells per well in 96-well plates 48 hours prior to infection in cell culture media (DMEM supplemented with 10% fetal bovine serum, 10% colony stimulating factor, 1% glutamax) and incubated at 37C and 5% CO_2_. Prior to infection, *M. abscessus* was cultured in 7H9 to an OD ∼0.6. Cultures were pelleted at 2850*g* for 5 minutes and washed twice with 1x PBS. Suspensions were centrifuged at 58*g* to pellet clumped bacteria. The supernatant was collected and diluted into DMEM supplemented with 5% horse serum and 5% FBS to achieve a multiplicity of infection of 2. Bacterial suspensions were added onto cells and spun at 335*g* for 10 minutes. Infected cells were washed with PBS and incubated in cell culture media at 37C. When necessary, media changes were performed every 2 days. For bacterial enumeration, BMMs were plated in clear TC-treated 96-well plates (Corning). Wells were washed three times with PBS, lysed in 100μl 0.01% Triton X-100 for 15 minutes, and the resultant bacterial suspensions were serially diluted in PBS supplemented with 0.05% Tween-80 and plated.

For macrophage viability assays, BMMs were plated in Optical clear-bottom black 96-well plates (Nunc). CellTiter Glo 2.0 was added to each well in a 1:1 ratio, plates were sealed with plate sealing tape (Nunc), and plates were gently agitated for 5 minutes to promote mixing.

Luminescence was read using a Molecular Devices SpectraMax M3 spectrophotometer (integration time: 500ms).

### In vivo infections

*M. abscessus* cultures were grown to an OD_600_ ∼0.6, pelleted at 2850*g* for 5 minutes, and washed twice with PBS. Suspensions were centrifuged at 58*g* to pellet clumped bacteria. The supernatant was collected and diluted in PBS to achieve the desired inoculum in a volume of 40 μl. Mice were anaesthetized with isoflurane, a pipette was used to instill 20 μl of inoculum into each nostril, and mice were monitored to ensure recovery. Mice were euthanized humanely, and lungs were harvested into 1ml of PBS 0.05% Tween-80. Lungs were mechanically homogenized and the resultant suspensions were serially diluted in PBS 0.05% Tween-80 and plated on 7H11 agar plates.

### Colony morphology

Cultures were grown to an OD_600_ ∼0.6, diluted, and plated on relevant media to an approximate density of 1-5 colonies per plate. Plates were incubated at 37C and imaged at 40x zoom on a Zeiss Axio Zoom V16 stereomicroscope with PlanNEOFLUAR Z 1.0x objective lens and AxioCam506 color camera.

### Ethidium bromide uptake

Membrane permeability to ethidium bromide was assayed as previously described (12). Cultures were washed in pre-warmed PBS containing 0.5% glucose to maintain cellular energization. Bacteria were resuspended to an OD_600_ of 0.5 in PBS 0.5% glucose 5μM ethidium, transferred to a black clear-bottomed 96-well plate, and ethidium fluorescence was monitored over 20 minutes using a Molecular Devices SpectraMax M3 spectrophotometer (excitation/emission: 520nm/595nm).

### Laurdan fluidity assay

Membrane fluidity was assayed via Laurdan as previously described (37). Cultures were grown in media lacking Tween-80 to an OD_600_∼0.6. Laurdan dissolved in dimethylformamide was added to a concentration of 10μM Laurdan and 1% v/v dimethylformamide and incubated at 37C with shaking for 2 hours. Cultures were pelleted at 2850*g*, washed several times in media supplemented with 1% v/v dimethylformamide, resuspended in 1/50^th^ culture volume, and transferred to a black clear-bottomed 96-well plate. Laurdan fluorescence was monitored using a Molecular Devices SpectraMax M3 spectrophotometer preheated to 37C (excitation/emission: 350nm/440nm and 350nm/490nm). The Laurdan generalized polarity was computed using the formula 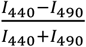, where I is the recorded fluorescence intensity at the specified wavelength.

### Detergent sensitivity assays

Cultures were grown to an OD_600_ ∼0.6, pelleted at 2850*g* for 5 minutes, and resuspended in fresh media. Cultures were then diluted to an OD_600_ of .05 in media containing detergent in 24-well plates, incubated for 24 hours at 37C with shaking, and the final OD_600_ was read using a Molecular Devices SpectraMax M3 spectrophotometer.

### Lipoarabinomannan detection

Cultures were grown in 7H9 lacking Tween-80 to an OD_600_ ∼0.6 and pelleted at 2850*g* for 5 minutes. Culture supernatant was filter sterilized twice through a 0.2μm Supor membrane (Pall Corporation). To assess cell pellet content, cells were resuspended in 1ml PBS, transferred to lysing matrix B tubes (MP Bio), and lysed using a [machine], resting on ice for 2 minutes between rounds. Debris was pelleted via centrifugation at 21130*g* for 15 minutes at 4C and the supernatant was filter sterilized twice through a 0.2μm Supor membrane. For western blot, cell pellet or culture supernatant fractions were standardized to culture OD_600_, boiled for 5 minutes in Laemmli buffer, and run on 4-20% Tris-Glycine criterion stain-free gel. The gel was imaged using stain-free gel fluorescence to confirm equivalent loading between lanes and transferred to a PVDF membrane overnight. Membranes were blocked for 1 hour with 5% milk in TBS-T. Membranes were incubated with CS-35 antibody conjugated to HRP (Creative BioLabs) in TBS-T 5% milk for one hour, washed several times in TBS-T, and visualized using the Opti-4CN colorimetric detection kit (BioRad).

For flow cytometry, pellets were fixed in formalin for 30 minutes and transferred to a V-bottom 96-well plate. All centrifugation steps occurred at 2850*g* for 5 minutes. Cells were washed several times with PBS, then blocked for 1 hour in PBS 5% bovine serum albumin fraction V (BSA). For primary staining, cells were incubated for 1 hour with CS-35 antibody (Creative Biolabs) in PBS 5% BSA and then washed several times in PBS 5% BSA. For secondary staining, cells were incubated for 1 hour with goat anti-mouse antibody conjugated to AlexaFluor 647, AlexaFluor 594, or AlexaFluor 488 (Invitrogen) in PBS 5% BSA. Cells were washed several times in PBS and measured on a BD Symphony A3. Data were analyzed in FlowJo.

### Hypoxia model

Cultures were grown in 7H9 lacking Tween-80 to an OD_600_ ∼0.6 and pelleted at 2850*g* for 5 minutes. Pellets were resuspended to an OD_600_ of 0.5 in 7H9 lacking Tween-80 and diluted in 7H9 lacking Tween-80 containing 45μg/ml methylene blue to 3*10^8^ cells in a final volume of 20ml. Cultures were transferred to flat bottomed glass tubes, stir bars were added, and tubes were sealed using rubber stoppers. Cultures were incubated at 37C with stirring at 180 rpm and oxygenation state was monitored through decolorization of methylene blue.

### Lipid extraction and analysis

Mycomembrane lipids were extracted as previously described (4). Briefly, cultures were centrifuged at 2850*g* for 5 minutes, supernatant was removed, and pellets were incubated with 10 mM dioctylsulfosuccinate in heptane overnight. Suspensions were centrifuged at 2850*g* for 5 minutes and supernatants were removed, dried under a gentle stream of nitrogen, and resuspended in chloroform.

Glycopeptidolipids were extracted as previously described (19). Briefly, cultures were centrifuged at 2850*g* for 5 minutes, supernatants were removed, and pellets were serially extracted with chloroform/methanol/0.3%NaCl (9:10:3 and 5:10:4 v/v/v, respectively). Extracts were combined, mixed with chloroform/0.3%NaCl (1:1 v/v) for 5 minutes, and centrifuged at 2850*g* for 10 minutes to separate aqueous and organic phases. Organic phases were removed, dried under a gentle stream of nitrogen, and resuspended in chloroform/methanol (2:1 v/v).

All extracts were analyzed using aluminum-backed silica gel plates (Silicycle). GPLs were separated using chloroform/methanol/water (90:10:1 v/v/v) (19). PIMs were separated using chloroform/methanol/13M ammonia/1M ammonium acetate/water (180:140:9:9:23 v/v/v/v/v) (4). TAGs were separated using toluene:acetone (99:1 v/v) (39). All TLCs were dipped in 50mM potassium permanganate 15mM sodium hydroxide 360mM sodium carbonate, dried, and gently heated for visualization.

## Data Availability

All data is available in the manuscript.

## Acknowledgements

We thank Jessica Seeliger (Stony Brook University) for helpful discussions and advice, and Varun Sridhar (University of California Berkeley) and Matt Traxler (University of California Berkeley) for technical assistance. We thank the Stanley lab for insights and advice. This work was funded by the NIH grant 1R01AI143722 to SAS.

---

**Supplementary Figure 1.**
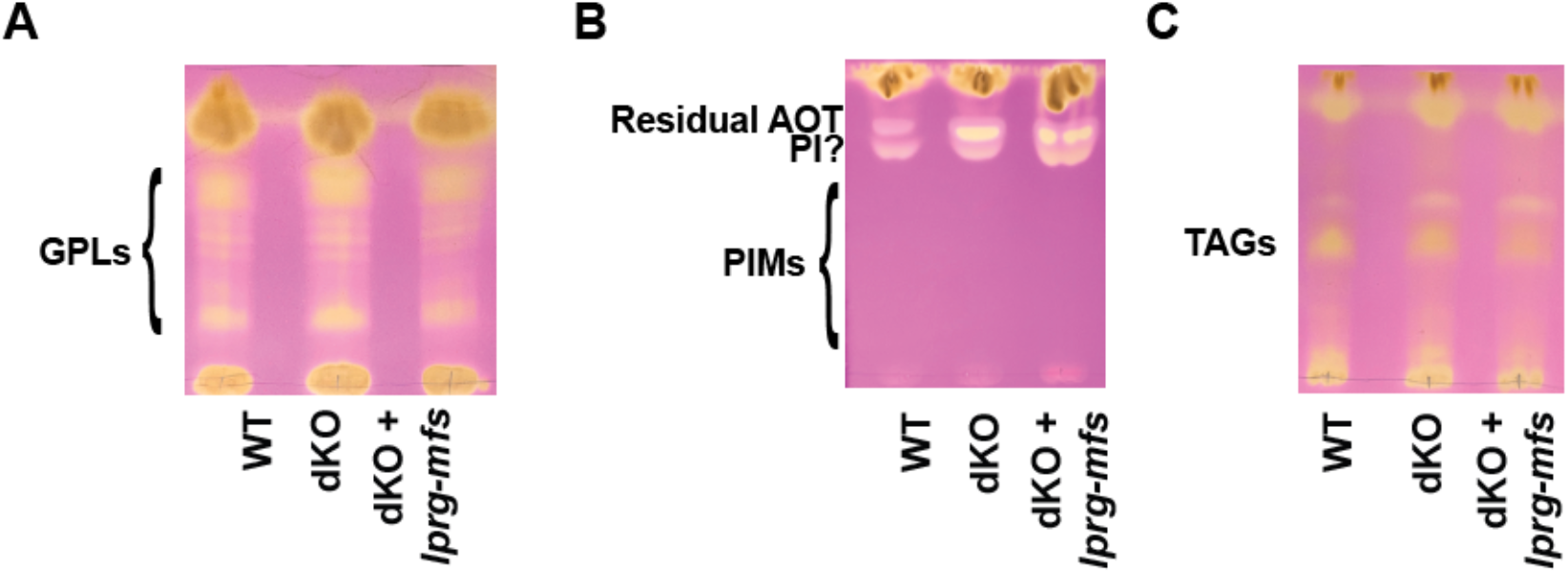
dKO *M. abscessus* does not have obvious changes in lipid abundance. **(A)** GPLs were extracted from cell pellets and analyzed by TLC using chloroform/methanol/water (90:10:1 v/v/v). **B**,**C:** Mycomembrane lipids were extracted via reverse micelle extraction and analyzed by TLC using chloroform/methanol/13M ammonia/1M ammonium acetate/water (180:140:9:9:23 v/v/v/v/v) **(B)** or toluene/acetone (99:1 v/v) **(C)**.

